# Separating error from bias: A new framework for facial age estimation in humans and AIs

**DOI:** 10.1101/2025.08.28.672503

**Authors:** Tzvi Ganel, Yarden Mazuz, Daniel Algom, Melvyn A. Goodale

**Author notes:** Corresponding author: Tzvi Ganel, Department of Psychology, Ben-Gurion University of the Negev, Beer-Sheva 84105, Israel.

## Abstract

Apparent facial age plays an important role in social interactions, serving a meaningful marker of biological aging. Although both humans and AIs achieve reasonable accuracy in estimating age from a person’s face, performance remains imprecise, leaving substantial room for errors and biases. Drawing on principles from classical psychophysics, we demonstrate that the existing literature on age estimation suffers from a critical theoretical and methodological shortcoming, which casts doubt on established findings. We show that the conventional measure used to benchmark the accuracy of human and AI performance is fundamentally confounded by response bias. Consequently, we introduce a novel measure that eliminates this confound. A revised framework based on simulated data, reanalysis of existing data, and new experimental results, reveals fresh insights into how facial age is processed by humans and AIs. Our structure opens up new directions for future research and applications in the study of aging.

## Introduction

Among the many different characteristics that people readily infer from a person’s face, age is considered primary^1^. Accurate identification of a person’s age is crucial for determining the nature of the social interaction involving the person, the person’s perceived level of health, attractiveness, and competence; apparent age serves as a marker of biological aging^2-4^. A youthful face is associated with positive characteristics such as increased attractiveness and health, which is probably the underlying force beyond the multibillion industry of anti-aging^5^. In the last decade, there has been a surge of interest in age estimation by automated AIs, which can be used for commercial and personalized medicine applications^4, 6^.

To evaluate the age of someone’s face, observers must consider a wide range of age cues, including the prominence of wrinkles and skin pigmentation, the overall shape of the face, and the person’s hairstyle and hair color, among others^7, 8^. Although these cues allow humans (and AIs) to achieve a considerable level of accuracy in evaluating age^6^, performance is still imprecise, limited by the available facial age cues, the observer’s perceptual resolution, top-down influences, as well as by genetic and environmental factors that cause people’s faces to age differently^2^. This leaves room for errors and biases that affect age evaluations from faces^9^.

Facial age cues dynamically change across the human lifespan^10-14^. From birth to adulthood, there are substantial changes in facial shape, which mainly originate from differences in the growth rates of different parts of the skull (e.g., cardioidal strain)^2^. The shape of the face changes to some degree even in adults^12^. Nevertheless, in adulthood, which is the main focus of the current study, the most dramatic age-related changes involve changes in the surface of the skin, including wrinkling, eye bags, age spots, as well as other features evident across the entire surface of the face^11^. It is not surprising, therefore, that the literature on age perception has focused on these surface-related changes in the appearance of adult faces. Amongst these cues, it has been suggested that wrinkling, especially in areas around the region of the eyes, is the most informative cue to someone’s age^9, 15^.

Cues such as wrinkling are typically less prominent in young compared to middle-aged and older adults, and, as a consequence, the mere existence of wrinkles in young adults can lead to them being perceived as older than they are^9^. In middle-aged and older adults, the accumulation of wrinkles as well as changes in skin pigmentation and sagging of the skin, are effective age cues^16^. Furthermore, in old adults, these surface-based cues are often accompanied by shape-related cues, such as the relative size of the ears and nose, providing a rich source of age-related information that could potentially lead to increased accuracy in age estimation with age. Beyond contributing to our understanding of how these different cues affect the psychometrics of age perception over the lifespan, careful measurement of their individual contributions as well as their interactions could provide insights into the underlying processing of facial age.

### A new framework for age estimation

#### Basic concepts in aging research

Unfortunately, conceptual and practical limitations in current research on age perception prevent the application of precise measurements of the accuracy of age estimation. The limitations cast doubts on the validity of several findings reported in the literature. Here, using insights from classic psychophysics (tailored to the domain of aging), we aim to identify and resolve these limitations. To do so, let’s first consider several key concepts with respect to age. The first and most straightforward is *chronological* (the “real”) age of a person. The second, and the main focus of the current paper, is the *perceived*, or *apparent* facial age. Although *chronological* (“real”) age and *perceived* or *apparent* facial age are positively correlated, this correlation is imperfect^17^, partly due to errors and biases in age estimation. The correlation is also less than perfect due to what apparent facial age uniquely implies about the fundamental process of aging^18^. Many would consider apparent age simply as a cue to the “true” chronological age of a person, but evidence from aging research suggests otherwise. Indeed, recent evidence strongly suggests that apparent facial age is an effective standalone marker of the *biological age* of a person.

Biological age is defined as the “time dependent functional decline that affects most living organisms”^19, 20^. Unlike the age-related concepts discussed earlier, there is no clear consensus about which particular measure provides the best estimate of biological age, but existing candidates include molecular measures such as DNA telomere length and methylation-based biomarkers^21^, as well as other measures such as those extracted from brain images^22^ and images of the retinal fundus and tongue^18^, and most relevant to the current study, from the visual appearance of faces^23^. There is growing evidence that apparent facial age (acquired from age evaluations of face photos either by human observers or by AIs) is an effective biomarker of aging^4, 17, 18, 21, 23-25^. For example, apparent age shows a higher correlation than chronological age with different biomarkers of biological aging^17^, an idea that receives additional support from a study of identical twins, showing that the differences in the estimated age from their photos was significantly correlated with biological markers of aging including mortality^25^. In addition, Belsky and colleagues^21^ acquired a new DNA biomarker of aging from a large group of participants of the same age cohort. The results showed that this biomarker was significantly correlated with perceived facial age. Recently, Bontempi and colleagues showed that apparent facial age (acquired from face photos by AI) is a significant marker for biological aging, longevity, and disease in cancer patients^4^. These findings highlight its potential application as an effective, low-cost biological marker for use in personalized medicine^26^, beyond the critical role of the perceived age of someone’s face in social interactions.

#### Errors in age estimation

Classical psychophysics has provided principles and empirical methods for studying human perception across many domains. Unfortunately, these principles have been insufficiently applied to the field of age perception.

In the most commonly used method for age estimation (both by humans and by AIs), the participant is asked to estimate a person’s age in years, typically from a photograph of the person’s face^3^.The main advantage of this method lies in its transparency; people readily extract the age of faces they see, which makes this task natural and straightforward. In a typical experiment in age perception, participants are presented with a series of face photos and are asked to estimate the age of the person in each photo in years^3, 6, 9, 27^. To avoid unwarranted effects of top-down influences and expectations, repeated presentations of photos of the same person (taken at the same age with different expressions, for example) to the same observer should be avoided^9, 15, 28, 29^. The *average perceived age* of each photo can then be calculated. In classical psychophysics, this measure parallels the concept of the Point of Subjective Equality (PSE), the perceived magnitude of a stimulus^30-32^. Note, however, that unlike many other psychophysical domains, such as stimulus size, where it is assumed that participants perceive size in a relatively similar manner across different types of stimuli, age perception is unique in that it also emphasizes individual differences between faces in the array of cues that signal age – and the saliency of those cues at different ages. Moreover, because people age differently, it is important to focus on potential differences along the perceived age of different individuals beyond the general facial attributes, such as their age group. To measure differences between faces, analyses of age estimation should focus on differences between faces (item-based analyses), rather than on traditional subject-based analysis. For example, an effective study of age perception should be tailored to identify both individual differences between people of the same chronological age (e.g., Joe is perceived as 3 years older than Jacob) as well as general group differences (e.g., young adults, on average, are perceived as 4 years older than their chronological age).

#### Age estimation bias

There are two types of errors in age estimation. The first is *bias*, defined as the *signed difference* between the average perceived age of a person and the person’s chronological age. For example, if Joe is 34 years old but is perceived on average to be 37, then bias is +3 years. In classical psychophysics, this difference between the PSE and the physical magnitude of the stimulus is termed the “constant error^33^. The term implies that bias should be relatively invariant across the range of stimulus magnitudes, which might be true for many different stimulus modalities such as brightness and size^34^, but is certainly not true for age. Studies from our lab and other labs have consistently demonstrated that bias in age estimation varies as a function of the chronological age of the person^3, 9^. In particular, faces of young adults are perceived, on average, as about 4 years older than their chronological age, whereas the faces of old adults are perceived to be about 5 years younger than their chronological age. This reversal of bias is robust and has been found across different facial databases and studies both in humans and in AIs^3, 6, 9, 35, 36^. Yet, its theoretical and practical significance has been largely overlooked. Here, we term this phenomenon as the Chronological Age Bias effect (CAB). The CAB has been suggested to result from people’s (and AI’s) tendency to use the estimated mean of a given distribution as the reference point for making judgments when uncertainty is a factor, i.e., *a regression to the mean effect*^3, 6^.

Additional sources of influence on biases in age estimations are gender and the emotional expression of the face. Previous studies have shown that female faces show smaller biases than male faces, and that this difference is modulated by the age of the face^3^. As well, studies from our lab and other labs have shown that, contrary to common belief, smiling faces are perceived as older than the faces of the same people when they have a neutral expression^15, 28^. This “Aging Effect of Smiling” (AES) is presumed to be driven by the formation of smile-related wrinkles in the region of the eyes. More recent data show that the AES is reduced with the chronological age of the face, is present for upright and inverted faces, for own and other-race faces, and with both human observers and AIs^6, 29, 37^. Beyond such group effects, however, there are individual differences between the faces of different people within each group that deserve careful attention. Different people appear to look younger or older compared to their peers (from the same age cohort). To identify these individual differences, group-dependent influences, which have been largely overlooked in previous literature, must be separated from biases that are idiosyncratic to the individual. Notably, the component of idiosyncratic age bias may be the one most closely related to the concept of Constant Error in psychophysics and is also a marker of one’s biological aging^19^. Note that in order to study age estimation effectively, a rigorous methodological design (a representative set of faces across the lifespan without repeated presentation of the same faces) should be accompanied by a comprehensive analysis that includes an item-based analysis of specific faces and ages. Such a design can then be used to compute the second type of error in age estimations, that of *absolute error*.

#### Absolute accuracy in age estimation

The second type of error in age estimation (as well as in virtually all other domains of human and machine estimation) refers to the absolute error of the response. Unlike the concept of bias, which has been theoretically underdeveloped, although correctly (statistically) measured, the accuracy of age estimation has been incorrectly approached both in theory and in practice. The accuracy of the response is linked to the concept of Just Noticeable Difference (JND), which refers to the perceptual resolution age. The JND assesses how accurately people (or AIs) can determine the magnitude (e.g., the apparent age) of a given stimulus. The larger the JND, the poorer is performance. Unlike bias, which is the signed error with respect to the *physical* magnitude of stimulus, the JND is measured independently from physical magnitude (and bias), in reference to the *perceived magnitude* of the stimulus (the PSE). For example, in the domain of size perception, which uses similar methodology to that used in age perception (i.e., estimations of the physical size of the object), the bias is the signed difference between the PSE and the physical magnitude, whereas the JND is most often measured by the (unsigned) variability of the response around perceived size (PSE)^38-40^.

Suppose that the person’s chronological age is 45 years, the physical magnitude of the stimulus. If the person’s age is judged, on average, to be 41 years, the bias is -4 years. Now consider the distribution of the judgments by different observers around the average of 41. Based on this distribution, a measure of dispersion or absolute accuracy can be calculated. The JND is such a measure. It is at this point that it becomes vital to recognize that the JND, or the absolute error of the age estimate, is calculated with respect to the mean of the *judgment*s, i.e., with respect to the average *apparent age*, not with respect to the true or chronological age. It becomes equally clear that the two measures – bias and absolute error – are independent. If bias is constant, then the correlation between a constant value and any and all other variables is zero. If bias varies, it is still uncorrelated with absolute error because the former refers to the physical magnitude, and the latter to perceived or estimated magnitude. Therefore, bias and absolute error represent two different types of inaccuracies and should be considered and measured independently from one another (for a similar idea in machine learning, see^41^). Ironically, the standard measure of error used in human and AI age estimation literature is essentially *inaccurate*, confounded with bias, which prevents its precise computation.

The current standard measure for absolute error in age estimation is the Mean Absolute Error (MAE). The MAE, a measure of the distribution, or variance, of the response around chronological age is computed by subtracting each response (age evaluation of a person’s face in years) from the chronological age of the face, and then by averaging the absolute values of these subtractions. Larger MAEs are associated with reduced accuracy. Because MAEs are computed with respect to the *chronological age* of the face rather than its *apparent age* (PSE), they are inevitably confounded with bias in nonlinear manner (with the only exception being a rare case in which perceived age exactly equals the chronological age). Moreover, due to the denser distribution of responses around the mean, MAEs will be erroneously inflated with larger group and individual biases (regardless of their direction). This idea is illustrated in Figure 1, which presents simulated data of item-based analysis in age estimations for faces of different chronological and perceived ages. Using the traditional, *confounded* measure of MAE leads to cases in which faces (or groups of faces) with *similar* (actual) degrees of accuracy are erroneously computed as being *different* in accuracy (Fig. 1, top panel). This could also lead to situations in which faces with *different* degrees of accuracy in the estimation of their apparent age are erroneously classified as having the *same* degree of accuracy (Fig. 1, bottom panel).

**Figure 1:**
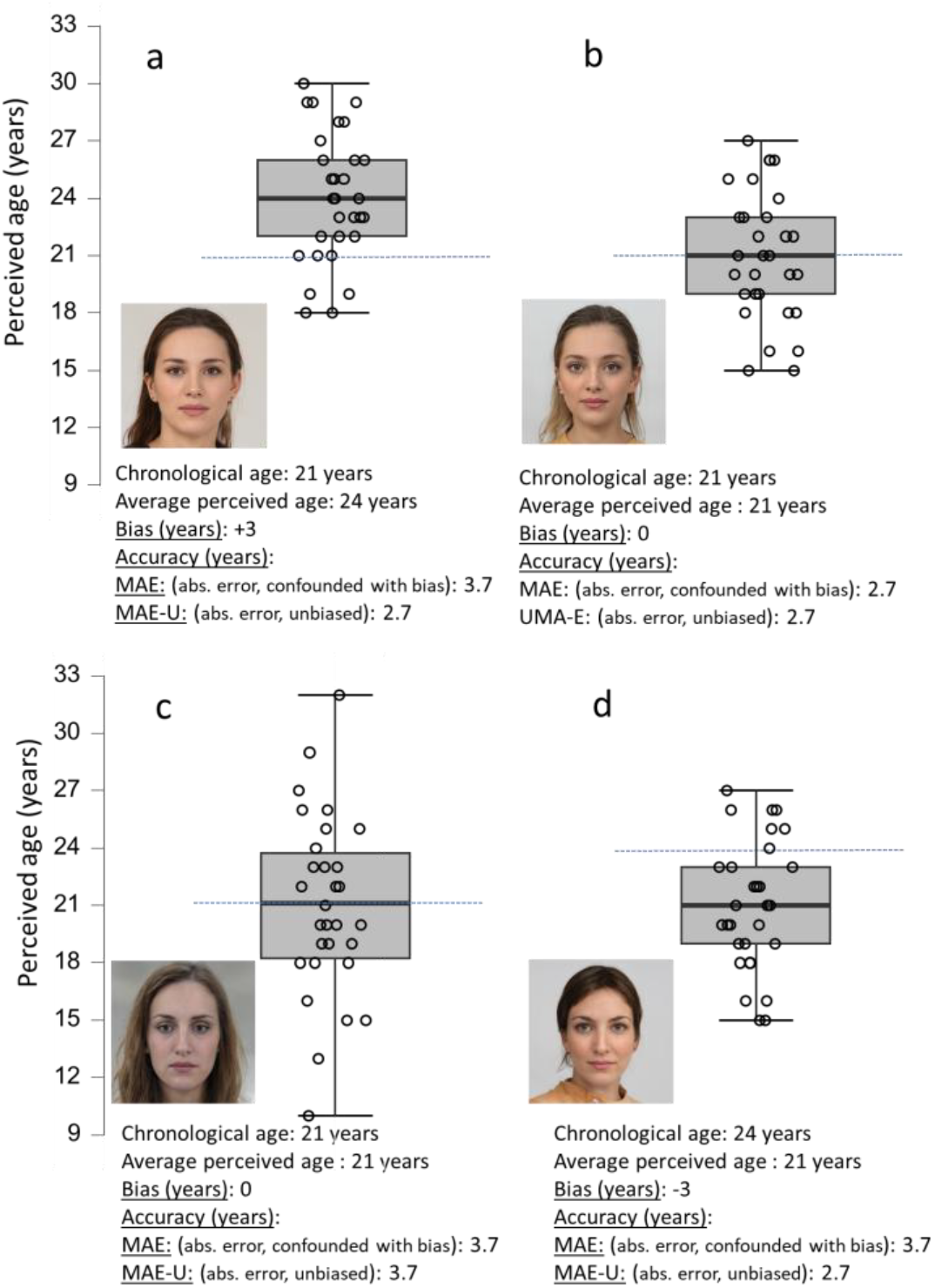
An illustration of the different types of errors in age estimation. The figure presents simulated data of age estimations of 30 “participants” for photos of 4 women (each dot represents a single response). There are two types of errors in age estimation: The bias (the average signed difference between the chronological age, represented by dashed lines, and the perceived age) and the absolute error (the variance of the response). The figure illustrates the idea that the most commonly used measure for accuracy - the MAE (mean absolute error, a measure of the distribution of the responses around the *chronological* age) is confounded with the bias of the response; MAEs are artificially inflated for faces (or group of faces) with larger biases. The MAE-U (mean absolute error, *unbiased*), on the other hand, for which absolute errors are computed around the average *perceived* age, is not confounded with bias. The inherent confound in MAEs could lead to errors in measuring accuracy. For example, faces *a* and *b* (top panel) are estimated with the same degree of accuracy around the perceived age (i.e., equal variance), yet their MAEs show an erroneous pattern of difference in accuracy due to confounds with biases. Faces *c* and *d* (bottom panel) are estimated with different degrees of accuracy around the perceived age, yet their MAEs appear to show a similar pattern of performance. Note, MAE-Us provide a unconfounded measure of performance in all cases. Note as well that with cases in which chronological and average perceived age are identical are extremely rare and are used here only for illustration. In most cases, there is at least some discrepancy (bias) between chronological and perceived age. These discrepancies are considerably larger (on average) for faces of old adults and for faces of young adults (the CAB effect, see Figure 2). The data in the figure illustrate the idea that the magnitude of bias (in any direction) erroneously inflates MAE values but has no effect on MAE-Us. Face images were computer-generated by Face-generator (https://generated.photos/face-generator).

In short, to measure absolute error of age estimation in an unbiased manner, the distribution or variability of responses should be computed with *reference to perceived, not to chronological, age*. This is done by computing the average absolute differences between each age evaluation and the *average perceived age of a face*. The result is an unconfounded measure of accuracy (in years) of age estimation. We term this error the Mean Absolute Unbiased error (MAE-U). As can be seen in the simulated data presented in Figure 1, MAE-Us (unlike the traditional, confounded MAEs) provide an unconfounded measure of accuracy.

Please note that due to the unique nature of the dimension of facial age, compared to other, more standard psychophysical dimensions (such as size, weight or brightness), absolute errors (that parallel the JND of the response) and biases are primarily based on an item rather than on the traditional subject-based analysis. For example, in a typical psychophysical study, the same participant is presented with many repetitions of the same stimulus, and the data are then used to compute the bias and the JND, separately for each participant (the bias is computed by the average signed difference between the participant’s responses and the physical magnitude and the JND is indicated by the variability^38^ of those responses around the perceived magnitude). Such a design would be improper for measuring individual differences between faces, which is a primary focus in age estimation research. Also recall that, due to possible top-down influences^9, 42, 43^, it is problematic to repeat the same face more than once to the same participant for age estimation. Therefore, an adapted, item-based analysis should be used to measure absolute errors and biases in age estimation. For this analysis, single responses of *different participants* to the *same item* are used as a basis to compute the absolute error (MAE-U) and the bias for each item. The MAE-U is computed based on the average absolute difference between each response and the mean perceived age; the bias is computed based on the average signed difference between each response and the chronological age (see Figure 1).

Now, having established a revised theoretical and practical framework for aging, we turn to using it to determine the true pattern of errors and biases in facial age estimation, both in humans and in AIs. Given that accuracy in age estimation was previously incorrectly measured, the current study is focused first on a critical re-evaluation of the pattern of errors and biases across the lifespan of adults faces (Study 1). To anticipate the main results of this first study, we show that the pattern of relation between chronological age and error across the adult lifespan is essentially different from what has been previously assumed, and that the processing of facial age varies with chronological age. In Study 2, we use insights from our first study to re-evaluate existing knowledge on differences in age estimation errors between humans and current AI technology. The new analysis reveals marked differences between humans and AIs pattern of performance, which dynamically change across the lifespan.

#### Study 1 – age estimation across the lifespan

Previous literature that looked at accuracy in age estimation throughout the adult lifespan found mixed results, probably due to the use of the MAE as the primary measure of error. As detailed before, the MAE is confounded with bias. Given that biases in age estimations change throughout the lifespan (e.g., CAB), it is likely that MAEs in age estimation throughout the lifespan do not represent the actual accuracy of the response. Given that the biases in age estimation for faces of middle-aged adults are smaller than the biases for the faces of old and young adults^9^, standard accuracy values were disproportionally inflated for faces of older and young adults compared to middle-aged adults. Furthermore, different studies used different face databases, which could vary with respect to individual biases and to the average biases of different age groups. Such variations could also have contributed to inconsistencies among the findings.

More than a decade ago, Voelkle and his colleagues published one of the most comprehensive studies of age perception^3^. 154 participants from different age groups evaluated the ages of 171 photos of the faces of young, middle-aged, and old adults, each presented with 5 different expressions. Relevant to the current discussion, MAEs were used as a (confounded) measure of accuracy, and the design included repeated presentation of faces of the same people with different expressions. The results showed that the errors increased with the age group of the target faces. On the face of it, this finding makes sense, if facial age cues become increasingly divergent with aging. As we discussed before, however, it is also possible that with increasing age, the accumulation of age cues may lead to the opposite effect, namely improving rather than decreasing performance with older faces. Critically, given that error has been always confounded with bias in previous studies (including studies from our lab) and given the large variation in these biases across the lifespan, it is difficult to draw a solid conclusion as to the relationship between chronological age and error. Other studies that have looked at age estimation along the lifespan also used MEAs as their primary measure ^44^. In most studies, there was again a general increase in error from young to old adulthood, but the rate of this increase differed substantially between studies^3, 44, 45^. In two more recent studies from our lab, in which we also used the confounded measure of MAE, there was again a general trend of an increase in error with age, although this relationship was not found consistently in all experiments and across all experimental conditions^9, 27^.

So, what is the true nature of the trajectory of errors across the adult lifespan? Do errors increase with age, as is currently assumed, or does this finding reflect a measurement confound? In Experiments 1a and 1b, we implemented our new framework to test the relation between error and chronological age in an extended and bias-free manner. We also used our framework to compare the traditional, confounded measure of error (MAE) to the newly developed measure (MAE-U). Finally, we used our framework to carefully examine the pattern of biases for the same set of faces across the adult lifespan (CAB). Our overarching aim is to provide a unified framework of facial aging research that can be implemented as a core model across the entire field. But first, let us establish the pattern of the relation between errors and chronological age across the adult lifespan.

### Experiments 1a and 1b

#### Methods

##### Participants

In Experiment 1a, we reanalyzed the data of Ganel and Goodale^9^ (Experiment 1a). Thirty students (18 females, mean age = 23.4 years, SD = 1.6 years) from Ben Gurion University of the Negev participated in this experiment. The participants in Experiment 1b were recruited online using the Prolific platform (75 participants, 36 females, mean age = 29.w years, SD = 10.7 years). The data of 5 participants, for which the mean absolute errors (MAEs) were substantially larger (more than 3 SD) from the group average were removed from the analysis. The experimental protocol was approved by the ethics committee of the Department of Psychology in Ben-Gurion University of the Negev. The study adhered to the ethical standards of the Declaration of Helsinki. All participants signed an informed consent form prior to their participation in the experiment. The manuscript contains no information or images that could lead to identification of a study participant.

##### Design and materials

The facial database in Experiment 1a was similar to one used in several of our previous studies^6, 9, 27^, with the exception that for the purpose of the present study, only the results of faces with neutral expression were analyzed. The database contains faces of 240 Caucasian individuals, equally divided to 3 age groups (40 females and 40 males in each group): young adults (20-40 years), middle-aged adults (40-60), and old adults (60-80 years). One potential issue with this set (as well as virtually all other sets used in the literature) was that the facial age of the photos is not equally divided to represent the entire range of chronological ages within each age group. This could limit the sensitivity to detect changes across the lifespan. To deal with this issue, and to test whether the findings extend to other faces and across different facial databases, we collected a new, extended and balanced set of 142 adult Caucasian face photos to be used in the present study (Experiment 1b) and in future experiments. 114 faces were taken from the Pal database^46^ and 28 from the UTKFace dataset^47^. Faces were cropped to the dimensions of 480X640 pixels, all with a neutral expression. None of the faces in the new set appeared in the first database.

Importantly, faces were equally distributed across the adult lifespan, with photos of one male and one female representing each chronological year within the age range of 18-89. Due to a technical error, one of the photos in Experiment 1b was presented twice and the data of this photo was therefore removed from the analysis.

##### Experimental procedure

The faces were presented in a random order. Each face was presented on the screen until a response – estimated age in years -- was made. Participants typed their response of years, which appeared below the target photo, and then pressed the “Continue” button to proceed to the next trial. To avoid top-down influences on age judgments, each face was presented only once throughout the experiment for each of the participants.

##### Data analysis

For each photo, we computed the average perceived age (i.e., the mean of the estimated age responses), the bias (the signed difference between each response and the chronological age), the MAE-U (average of the non-signed absolute differences around perceived age – measured by the difference between each response and the average perceived age of a particular face), and the MAE (average of the absolute differences around chronological age). In the current study, data was analyzed in an item-based analysis, which allows a detailed exploration of the pattern of results across the lifespan. We note that the data analysis could also be extended to a traditional subject analysis, which does not allow detailed exploration of the pattern of results across the lifespan, providing only a general measure of performance for a condition/group of faces. The general pattern of results of the subject-based analysis was similar to the item-based analysis. For sake of brevity, we do not include this analysis in the current paper. Data analysis was performed using the R and JASP platforms.

## Results and discussion

The pattern of the errors and biases across the lifespan in experiments 1a and 1b is presented in Figure 2. The pattern of results is similar across the two experiments. First, the actual error rate (measured by the MAE-Us) is substantially smaller compared to the traditional, confounded error rate (measured by MAEs), due to the removal of the confounds from biases. Most importantly, and unlike what has been previously assumed, errors do not increase monotonically with chronological age. Instead, the absolute error increases with age up to mid-adulthood and then begins to decrease. These findings suggest (perhaps counterintuitively) that for age estimations of old faces, human observers become increasingly more, not less, accurate, for faces of old adults. This phenomenon could be accounted for by the accumulation of age-related cues in old adulthood, as well as by possible qualitative changes in the relative reliance on surface and on shape-related cues with different age groups of faces. Note as well that the results of Experiment 1b provide a more sensitive measure of the error trajectory across the lifespan compared to Experiment 1a (and previous studies), presumably due to the equal distribution of faces across the lifespan and the larger stimulus range of stimuli.

**Figure 2:**
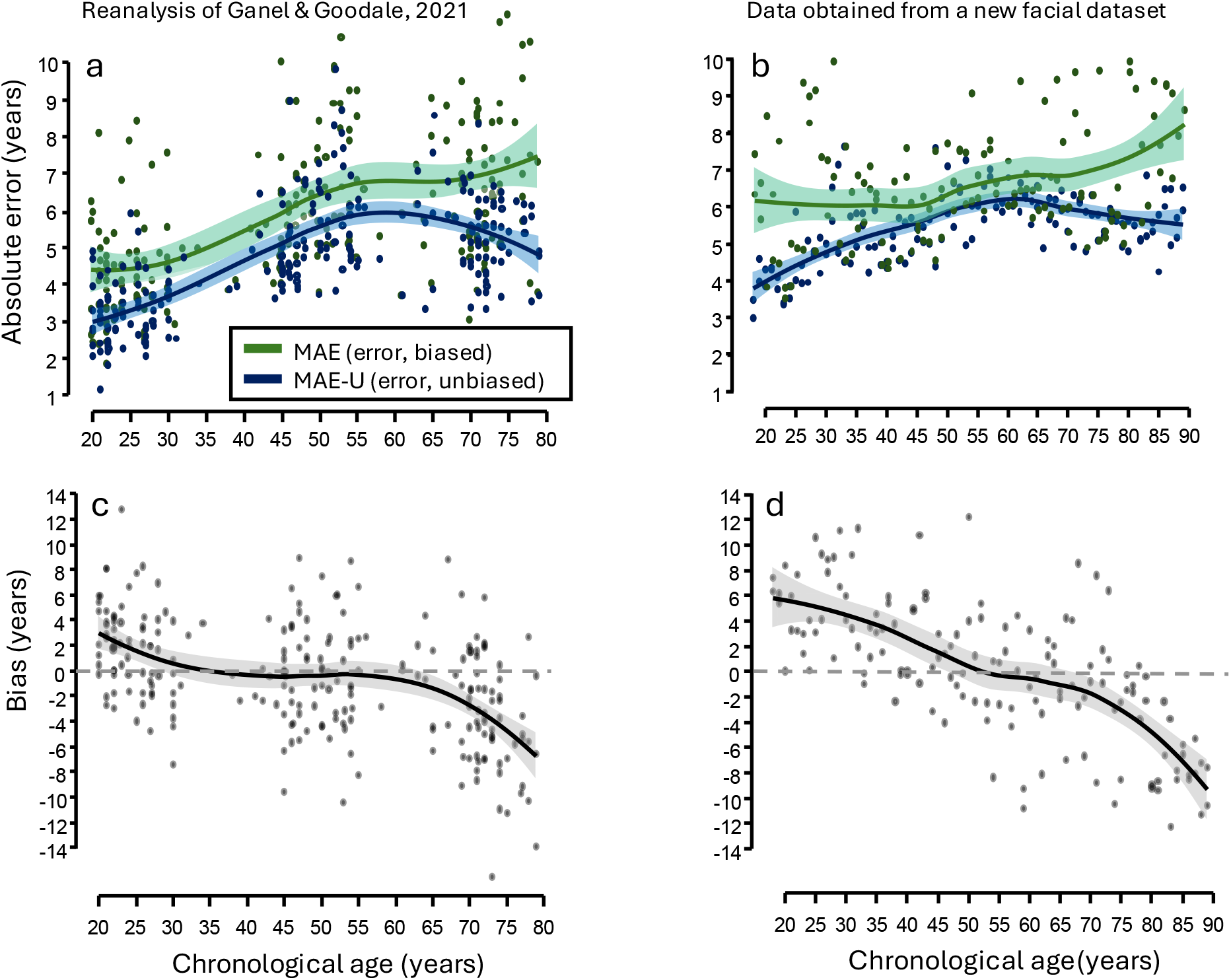
The trajectory of errors across the adult lifespan in experiments 1a and 1b. Two different facial databases were presented in the two experiments. The lines represent the best fit of the data across chronological age using the “loess” function in Flexplot addon (JASP). The top panels present the absolute errors throughout the lifespan. The green lines show the traditional, confounded measure (MAE). The blue lines show the newly developed, unbiased measure (MAE-U). Note the substantial difference along the pattern of results for the two measures. MAEs show a (false) trend of monotonic decrease in performance with age. This trend is confounded by large biases for faces of old and young adults compared to middle-aged adults (see bottom panels). When these confounds are removed (MAE-Us), the actual pattern of errors is revealed: Error increases with age from early to mid-adulthood, and then begins to decrease. Please also note that after the removal of confounds, the overall error rate is substantially smaller than what has previously been assumed.

To statistically compare the pattern of errors across the lifespan between MAE-Us and MAEs, we first performed a categorical analysis of the two error measures. To this end, we divided the range of chronological age into 6 specific age groups (20-29, 30-39, and so on) in Experiment 1a and to 7 specific age groups in Experiment 1b. A mixed ANOVA analysis was used to compare the pattern of results, with type of measure (MAE-Us vs MAEs) as the within-subject (item) independent variable and with the gender of the photo and specific age group as the between-subjects variables. Separate analyses were performed for Experiments 1a and 1b. The pattern of results was similar for the two experiments, so the results are reported in a unified fashion. For sake of brevity, only significant effects are reported.

A main effect of *measure type* (MAE-U vs MAE) was found in both experiments [exp 1a: F(1,228) = 63.7, p < 0.001; exp 1b: F(1,129) = 125.9, p < 0.001]. This indicates that the magnitude of errors was smaller in MAE-Us compared to MAEs, due to the removal of confounds inherent to biases. A main effect was also found for specific age group [exp 1a: F(5,228) = 32.3, p < 0.001; exp 1b: F(6,129) = 7.3, p < 0.001], indicating general differences in absolute errors between different age groups. More importantly, a significant interaction was found between *age group* and between *type of measure* [exp 1a: F(5,228) = 1.9, p < 0.5, one tailed; exp 1b: F(6,129) = 6.7, p < 0.001]. This finding provides the first indication that the pattern of errors throughout the lifespan is different between MAE-Us and MAEs. More specifically, as can be seen in Figure 2 (top panel), MAEs monotonically increased with age throughout the lifespan. In sharp contrast, the unbiased measure of MAE-U increases with age in a largely monotonic fashion up to mid-adulthood, but then begins to decrease. To further test if MAE-Us show such a decrease, we performed specific comparisons between the age group of 70-79 and the age group of 60-69. In both experiments, the results showed that MAE-Us were significantly smaller for the age group of 70-79 [exp 1a: t=2, p < 0.05; exp 1b: t=3.1, p < 0.05].

Regression models were used to further explore the nature of the relationship between chronological age and absolute errors, but now in a continuous fashion. First, we compared a linear versus a quadratic model as a fit for the MAE-U data across the lifespan. The results (Tables 1 and 3) clearly show that the data is best accounted for by a quadratic relation between MAE-U and age, in both Experiments 1a and 1b. The negative coefficient value for the quadratic fit indicates a curvilinear (inverted-U) fit for the relation between MAE-Us and chronological age in both experiments (see Figure 2). Unlike MAE-Us, the regression analysis of MAEs showed a different relationship between absolute error and chronological age. As can be seen in Tables 2 and 3, the MAE data is best accounted for by a linear rather than a quadratic fit with chronological age.

**Table 1:** Linear and quadratic regression models for MAE-Us (unbiased errors)

| **Linear model** |  |  |  |  |  |  |
| --- | --- | --- | --- | --- | --- | --- |
|  | Experiment 1a |  |  | Experiment 1b |  |  |
| *Predictors* | *Estimate* | *CI* | *p* | *Estimate* | *CI* | *p* |
| (Intercept) | 2.54 | 2.12 – 2.95 | <0.001 | 4.28 | 3.91 – 4.65 | <0.001 |
| age | 0.05 | 0.04 – 0.05 | <0.001 | 0.02 | 0.02 – 0.03 | <0.001 |
| **Linear + Quadratic model** |  |  |  |  |  |  |
| (Intercept) | 4.73 | 4.59 – 4.87 | <0.001 | 5.44 | 5.33 – 5.56 | <0.001 |
| age [linear] | 13.58 | 11.42 – 15.73 | <0.001 | 5.40 | 4.00 – 6.80 | <0.001 |
| age [quadratic] | -8.11 | -10.26 – -5.95 | <0.001 | -5.03 | -6.42 – -3.63 | <0.001 |

**Table 2:**
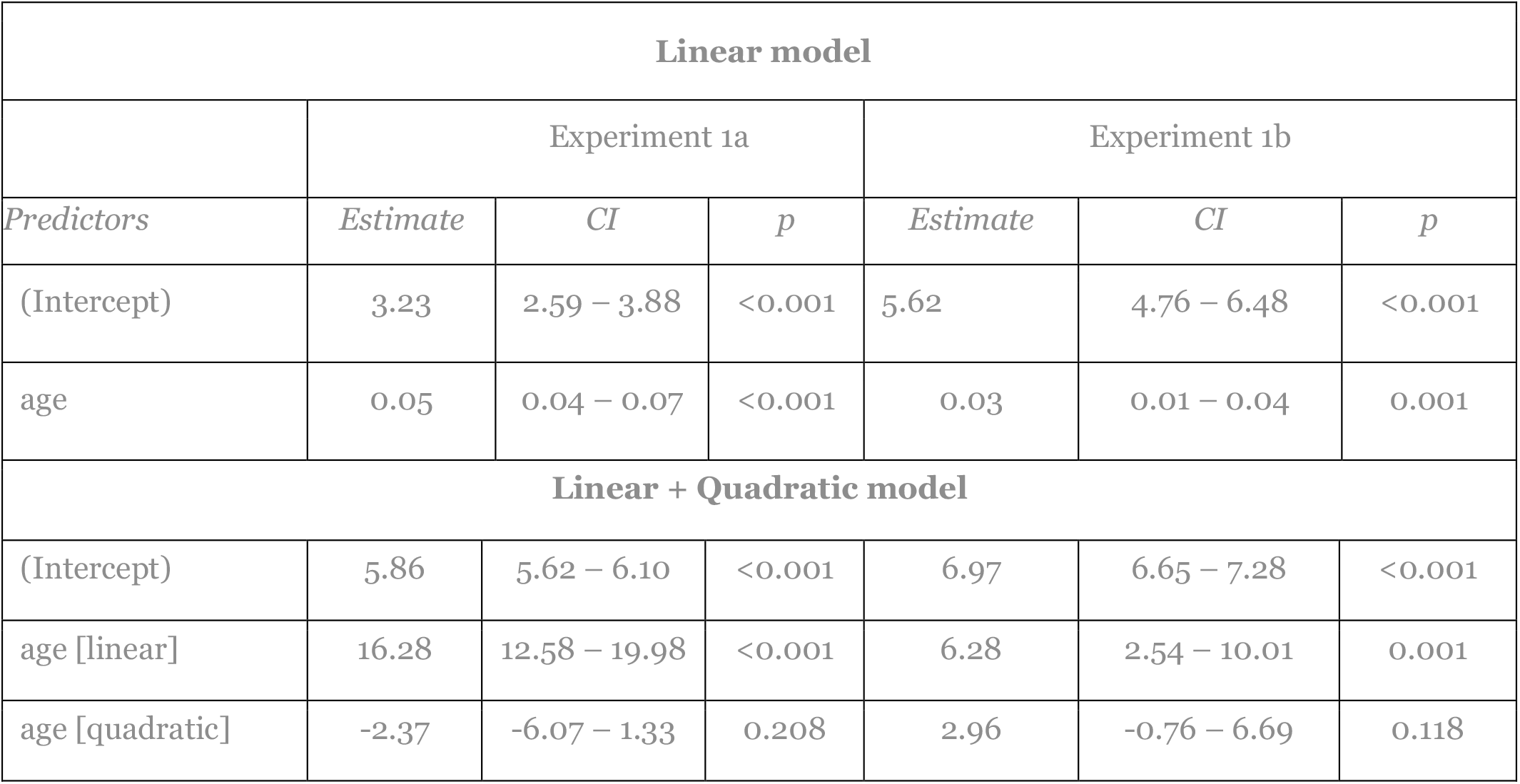
Linear and quadratic regression models for MAEs (biased errors)

| **Linear model** |  |  |  |  |  |  |
| --- | --- | --- | --- | --- | --- | --- |
|  | Experiment 1a |  |  | Experiment 1b |  |  |
| *Predictors* | *Estimate* | *CI* | *p* | *Estimate* | *CI* | *p* |
| (Intercept) | 3.23 | 2.59 – 3.88 | <0.001 | 5.62 | 4.76 – 6.48 | <0.001 |
| age | 0.05 | 0.04 – 0.07 | <0.001 | 0.03 | 0.01 – 0.04 | 0.001 |
| **Linear + Quadratic model** |  |  |  |  |  |  |
| (Intercept) | 5.86 | 5.62 – 6.10 | <0.001 | 6.97 | 6.65 – 7.28 | <0.001 |
| age [linear] | 16.28 | 12.58 – 19.98 | <0.001 | 6.28 | 2.54 – 10.01 | 0.001 |
| age [quadratic] | -2.37 | -6.07 – 1.33 | 0.208 | 2.96 | -0.76 – 6.69 | 0.118 |

**Table 3:** A comparison between the linear and the quadratic regression models for MAE-Us and MAEs

|  | MAE-Us |  |  |  |  |  |  |  |
| --- | --- | --- | --- | --- | --- | --- | --- | --- |
|  | Experiment 1a |  |  |  | Experiment 1b |  |  |  |
| Model | AIC | BIC | R<sup>2</sup> | Adj. R<sup>2</sup> | AIC | BIC | R<sup>2</sup> | Adj. R<sup>2</sup> |
| Linear | 776.88 | 787.32 | 0.35 | 0.34 | 352.8 | 361.69 | 0.24 | 0.23 |
| Linear + Quadratic | 728.78 | 742.70 | 0.47 | 0.46 | 310.66 | 322.51 | 0.44 | 0.43 |
|  | MAEs |  |  |  |  |  |  |  |
| Model | AIC | BIC | R<sup>2</sup> | Adj. R<sup>2</sup> | AIC | BIC | R<sup>2</sup> | Adj. R<sup>2</sup> |
| Linear | 988.2 | 998.65 | 0.24 | 0.24 | 591.75 | 600.64 | 0.07 | 0.06 |
| Linear + Quadratic | 988.59 | 1002.52 | 0.24 | 0.24 | 591.25 | 603.09 | 0.09 | 0.07 |

These results reinforce the idea that, unlike what has been previously assumed, errors in age estimation (computed by the unbiased measure of MAE-U) do not monotonically increase with age, but instead show a curvilinear relation, an increase up to a certain point (approximately at the age of 60 years), and then a decrease. We will discuss later possible accounts for mechanisms that could mediate this pattern of human performance. But first, we turn to an examination of the pattern of errors in the estimation of age by AIs. To this purpose, we used our framework in a second study to perform a comprehensive comparison between the pattern of errors and biases in humans and AIs.

### Study 2 – age estimations in humans and AIs

In this study, we used our newly developed framework to compare AI and human performance in age estimation. There is a huge interest in automated AI technology for identifying and estimating the age of individuals from their faces^4^. In what ways do AIs differ from humans in carrying out these tasks? In a study conducted several years ago, we compared human performance with the performance of a large sample (21) of the most prominent AI technology available at the date of the evaluation^6^. The AIs included Microsoft Face API, Amazon Rekognition, Everypixel API as well as other prominent AIs (see^6^, Table 1). The facial database used for humans and AIs was identical to the one used in Experiment 1a. However, the analysis in our original study did not include a comprehensive investigation of the pattern of errors and biases across the lifespan. More importantly, for the categorical analysis that was performed in the original study, errors were calculated using the confounded measure of MAE. The results showed marked similarities between human and AI performance in the pattern of errors and biases. Notably however, AI performance was more heavily affected by biases and inaccuracies than humans^6^. For example, AIs showed a more pronounced Chronological Age Bias (CAB) compared to humans, mainly due to larger biases (age underestimation) in faces of older adults. In addition, AIs showed larger aging effect of smiling (AES) compared to human observers.

More relevant to the current study, for both humans and AIs there was a monotonic increase in accuracy (measured by MAEs) with age group. Still, this decrease was significantly larger for AIs. Based on our newly developed framework for measuring errors and on the findings from Experiments 1a and 1b that show that errors (MAE-Us) do not monotonically increase across the entire lifespan (at least in humans), we reanalyzed the results of humans vs. AIs in a corrected and a comprehensive manner. First, we compared the error rates between the traditional, confounded measure of MAE and between the newly developed measure (MAE-U). In addition, and unlike what was done in previous studies, we used an item-based analysis to compare the pattern of errors and biases in humans and AIs across the adult lifespan. We asked the following questions: Is the true, unconfounded pattern of errors (MAE-Us) across the lifespan similar between humans and AIs? Is AI performance similar overall to human performance across the adult lifespan?

### Experiment 2

#### Methods

##### Participants

The data was taken from Ganel et al.^6^ in which we compared the performance of 21 commercial and non-commercial AI technology (for a complete list, see Table 1 in^6^) to human performance. The human participants were the same 30 BGU students, whose data were analyzed in Experiment 1a.

##### Design, procedures, and analysis

The facial database was identical for humans and AIs and was same as that used in Experiment 1a and in our previous studies^6^. The procedure was also identical to the one used in Experiment 1a. For sake of brevity, the current study focuses only on the faces with the neutral expression. For each photo, MAE-Us, MAEs, and biases were computed separately in participant group (humans and AIs).

##### Results and discussion

The results of the AI and human patterns of performance across the lifespan are presented in Figure 3. Note the marked differences between the pattern of performance indicated by the confounded measure of MAE and between the unconfounded measure of MAE-U. The traditional analysis of MAEs seems to show that humans and AIs have the same pattern of performance, with errors increasing with age. However, the unbiased pattern of performance is different between humans and AIs; unlike the errors in human age estimation, which show a curvilinear relation with age (also see Study 1), errors in AI show a monotonic increase with age across the lifespan. This dissociation along the pattern of absolute errors between humans and AIs is detected only when biases are effectively removed (MAE-Us). Also note, that across errors and biases, AIs show an advantage in age estimation performance for the faces of young and middle-aged adults, while humans outperform AI technology for faces of old adults.

**Figure 3:**
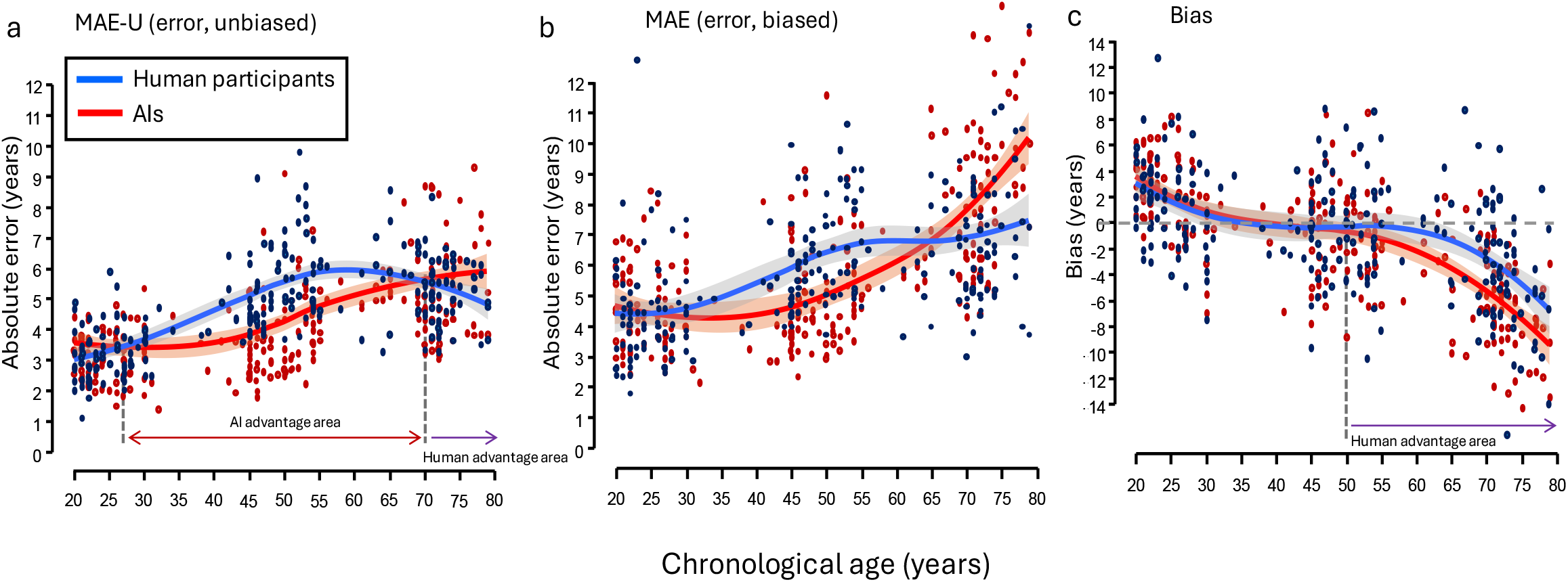
eanalysis of anel et al. (2022, Experiment 1a). The trajectory of human and AI performance across the adult lifespan. The lines represent the best fit of the data across chronological age using the “loess” function in Flexplot addon (JASP). When biases are effectively removed (a), a dissociation is found between human and AI pattern of performance. Unlike humans, AIs show a monotonic increase in errors with age (a). This dissociation was undetected when the confounded measure of MAEs was used (b). Note, that there is a larger Chronological Age Bias effect (CAB) in AI compared to human performance (c). Also note, that AIs generally outperform humans across the adult lifespan, with the exception of the faces of old adults, for which humans show an advantage, both for errors and for biases.

Given that group type (AI compared to Humans) served as a within-item factor, we used Hierarchical Linear Modeling (HLM) regression model to compare human and AI performance across chronological age in a continuous manner. The dependent measure was the absolute unbiased error indicated by MAE-Us. Based on the quadratic nature of the relation of MAE-Us with age established for human data in Study 1, we used this quadratic model to analyze the data. Table 4 shows the resulting MAE-Us using a quadratic mixed-effect modeling of chronological age, group type, and the interaction between the two. As can be seen in the table, the linear effect of age is positive and significant, suggesting that MAE-Us initially increased with age. The quadratic effect of age is negative, indicating that overall, the age-related increase slowed or reversed at older ages. A main effect of Group indicates that overall, AIs had smaller errors compared to humans. More importantly, the significant interaction between the quadratic component of age and group indicates that human errors are better accounted for by a curvilinear relation with age compared to errors in AI. This analysis reinforces the idea that the pattern of error in AIs is essentially different from that observed in humans.

**Table 4:** Quadratic mixed-effects model of MAE-Us predicted by group type and chronological age

| *Predictors* | *Estimate* | *CI* | *p* |
| --- | --- | --- | --- |
| (Intercept) | 4.73 | 4.57 – 4.89 | <0.001 |
| age [linear] | 13.58 | 11.42 – 15.73 | <0.001 |
| age [quadratic] | -8.11 | -10.26 – -5.95 | <0.001 |
| Group | -0.31 | -0.52 – -0.10 | 0.004 |
| age [linear] × Group | -0.09 | -4.67 – 4.49 | 0.969 |
| age [quadratic] × Group | 15.59 | 11.01 – 20.17 | <0.001 |
| $\sigma^2$ | 1.36 | | |
| $\tau_{00\ p}$ | 0.07 | | |
| ICC | 0.05 |  |  |
| $R^2$ / $R^2$ adjusted | 0.398 / 0.429 | | |

One possible explanation for different types of correspondence between chronological age and error in AI could be that, unlike human observers, who can effectively utilize accumulating age-related cues in old adulthood, AI technology is still lacking in this domain. This idea receives further support from the relatively large (negative) biases in AIs compared to humans for old adults faces (Figure 3c). Importantly, unlike in our original report and unlike in virtually all previous studies in the aging domain, the new analysis in Study 2 effectively distinguishes between error and bias, so the two types of inaccuracies can be independently explored.

It is interesting to speculate that the current focus on AI age estimation in personal medicine in older adults^4^ could lead to advances in AI technology for age estimation in this age group. It is likely that the commercial and entertainment-based focus on automated age estimation in young adults resulted an emphasis on training AI models on faces from this young age group and an absence of training on faces of older adults. This focus may now shift due to current work demonstrating the crucial role of apparent facial age as a biological marker for aging, particularly in old adulthood^4, 25, 26^. Our framework for age estimation provides a promising tool to follow up on possible advances in AI technology across the lifespan and we plan to implement it for this purpose in future studies.

##### General Discussion

Our primary aim was to provide a theoretical framework for understanding the estimation of age from faces that overcomes current limitations. Earlier investigations of the perception of age from faces have routinely confounded measures of error with measures of bias. We addressed this issue by providing a new, unbiased measure of absolute accuracy in age estimation.

In Studies 1 and 2, we demonstrate that once error is measured in a bias-free manner, new insights emerge with respect to how facial age is perceived across the adult lifespan. The results of Study 1 show that, unlike what has been previously assumed, accuracy in age estimation does not decrease in a linear or monotonic fashion with chronological age. Instead, across different facial stimuli and in different experiments, human accuracy decreases with faces from age 20 to about age 60 and then begins to increase. We speculate that this pattern of results reflects the ability of human observers to rely on cues to age that accumulate as people grow older.

Surprisingly, the results of Study 2 showed that the estimation of age by AIs does not show the same pattern of errors over the lifespan shown by human observers, presumably reflecting the use of truncated training sets and their lack of exposure to older adults.

We hope that our new framework will not only redress current problems in the field but will also enable new directions for research in age perception. One such direction is the role of the perceiver with respect to accuracy and bias in age estimation. In the current paper, we have focused mainly on the attributes of the facial stimuli and changes in their prominence and distribution over the lifespan. It is very likely, however, that attributes of the perceiver, such as their age^3^ and differences in their exposure to different age groups and their attention to the range of cues to age will affect their estimates. Our framework provides practical tools to tease apart biases from errors in age estimation to investigate, for example, how the age of the perceiver affects the estimation of age of faces across the lifespan.

A well-known example of such stimulus-observer interactions in age perception is referred to as the “own-age advantage”^36^. In general terms, an own-age advantage in age perception refers to the idea that a given age group should show better age estimation performance for faces of their own age group compared to the faces of other age groups. As detailed before, such improvement can be manifested in terms of bias or accuracy. For example, young adults are expected to show smaller errors in estimating the ages of young adults compared to middle-aged and older adults, while at the same time, older adults should show the inverse pattern of performance, doing better with older than younger faces. Surprisingly, a review of the literature shows that although the term “own-age advantage” is widely used, there is scarce evidence to support its presence. Studies that looked at possible own-age advantages in age estimations either found no evidence of own-age advantages, or partial evidence to support such effects^3, 16, 36, 44, 48, 49^.

This issue becomes even more complex given that all previous studies of the own-age advantage used a confounded measure for error. For example, in Voelkle et al.’s study, analysis of accuracy (MAEs) did not show any own-age advantage effect for errors and only a partial trend for own-age advantage for biases^3^. We hope that our framework will be used in future studies to effectively tease apart bias from error in order to look at potential interaction between the attributes of facial stimuli (e.g., their chronological age, gender, ethnicity, etc.) and the attributes of the perceiver (e.g., the perceiver’s age group, gender, ethnicity, and face identification skills).

It should be noted that, although our new framework reveals for the first time how error and bias can be disentangled in age estimation studies, most of the work in the field (including our own) has used photos of faces, where only static information is available. In future studies, the use of video clips would make it possible to explore the role of dynamic cues^50^, and their interactions. The use of videos would also make it possible to examine how differences in gait and other body movements across the lifespan contribute to our estimates of age and to expand the study of ageing to other perceptual domain beyond vision. For example, how people extract age information from voices^51^. Importantly, our framework will make it feasible to examine whether these additional sources of information impact the accuracy of age judgements across the lifespan differently from any biases that might be present. Finally, we suspect that the framework could also be usefully applied to other domains of perception where multiple cues contribute to the judgements being made, particularly when the prominence of those cues and their relative weightings vary across time (or other dimensions).

## Acknowledgement

We thank Yoav Kessler and Mattan Ben-Shachar on their advice on earlier drafts of this manuscript. Item analysis data is available online at https://osf.io/fzq89/.

## References

1. George, P. A. & Hole, G. J. Recognising the ageing face: the role of age in face processing. Perception 27, 1123–1124 (1998).

2. Rhodes, M. G. Age estimation of faces: A review. Applied Cognitive Psychology 23, 1–12 (2009).

3. Voelkle, M. C., Ebner, N. C., Lindenberger, U. & Riediger, M. Let me guess how old you are: effects of age, gender, and facial expression on perceptions of age. Psychol. Aging 27, 265–277 (2012).

4. Bontempi, D. et al. FaceAge, a deep learning system to estimate biological age from face photographs to improve prognostication: a model development and validation study. Lancet Digit. Health. 7, 100870 (2025).

5. Balasubramanian, S. Anti-aging and aesthetic medicine: the silent rise of this multibillion dollar industry. (2020).

6. Ganel, T., Sofer, C. & Goodale, M. A. Biases in human perception of facial age are present and more exaggerated in current AI technology. Sci. Rep. 12, 22519–w (2022).

7. Lai, M., Oruc, I. & Barton, J. J. S. The role of skin texture and facial shape in representations of age and identity. Cortex 49, 252–265 (2013).

8. George, P. A. & Hole, G. J. The role of spatial and surface cues in the age-processing of unfamiliar faces. Visual Cognition 7, 485–509 (2000).

9. Ganel, T. & Goodale, M. A. The effect of smiling on the perceived age of male and female faces across the lifespan. Sci. Rep. 11, 23020–2 (2021).

10. Gonzalez-Alvarez, J. & Sos-Pena, R. The role of facial skin tone and texture in the perception of age. Vision Res. 213, 108319 (2023).

11. Porcheron, A., Mauger, E. & Russell, R. Pmc3590275; Aspects of facial contrast decrease with age and are cues for age perception. PLoS One 8, e57985 (2013).

12. Mark, L. S. et al. Wrinkling and head shape as coordinated sources of age-level information. Percept. Psychophys. 27, 117–124 (1980).

13. Ramanathan, N. & Chellappa, R. Modeling age progression in young faces (2006 IEEE computer society conference on computer vision and pattern recognition (CVPR’06) Ser. 1, IEEE, 2006).

14. Mileva, M., Young, A. W., Jenkins, R. & Burton, A. M. Facial identity across the lifespan. Cognit. Psychol. 116, 101260 (2020).

15. Ganel, T. Smiling makes you look older. Psychonomic Bulletin and Review 22 (2015).

16. George, P. A. & Hole, G. J. Factors influencing the accuracy of age estimates of unfamiliar faces. Perception 24, 1059–1073 (1995).

17. Umeda-Kameyama, Y. et al. Cognitive function has a stronger correlation with perceived age than with chronological age. Geriatr. Gerontol. Int. 20, 779–784 (2020).

18. Wang, J. et al. Accurate estimation of biological age and its application in disease prediction using a multimodal image Transformer system. Proceedings of the National Academy of Sciences 121, e2308812120 (2024).

19. Lopez-Otin, C., Blasco, M. A., Partridge, L., Serrano, M. & Kroemer, G. The hallmarks of aging. Cell 153, 1194–1217 (2013).

20. Hastings, W. J., Almeida, D. M. & Shalev, I. Conceptual and Analytical Overlap Between Allostatic Load and Systemic Biological Aging Measures: Analyses From the National Survey of Midlife Development in the United States. J. Gerontol. A Biol. Sci. Med. Sci. 77, 1179–1188 (2022).

21. Belsky, D. W. et al. DunedinPACE, a DNA methylation biomarker of the pace of aging. Elife 11, 10.7554/eLife.73420 (2022).

22. Levakov, G., Rosenthal, G., Shelef, I., Raviv, T. R. & Avidan, G. From a deep learning model back to the brain—Identifying regional predictors and their relation to aging. Hum. Brain Mapp. 41, 3235–3252 (2020).

23. Avila, F. R. et al. Perceived Age as a Mortality and Comorbidity Predictor: A Systematic Review. Aesthetic Plast. Surg. 47, 442–454 (2023).

24. Dykiert, D. et al. Predicting mortality from human faces. Psychosom. Med. 74, 560–566 (2012).

25. Christensen, K. et al. Perceived age as clinically useful biomarker of ageing: cohort study. BMJ 339, b5262 (2009).

26. Meng, D., Zhang, S., Huang, Y., Mao, K. & Han, J. J. Application of AI in biological age prediction. Curr. Opin. Struct. Biol. 85, 102777 (2024).

27. Ganel, T. & Goodale, M. A. Smiling makes you look older, even when you wear a mask: the effect of face masks on age perception. Cogn. Res. Princ Implic 7, 84–3 (2022).

28. Ganel, T. & Goodale, M. A. The effects of smiling on perceived age defy belief. Psychonomic Bulletin and Review (2017).

29. Yoshimura, N. et al. PMC7797192; Age of smile: a cross-cultural replication report of Ganel and Goodale (2018). J Cult Cogn Sci, 1–15 (2021).

30. Marks, L. E. & Gescheider, G. A. in 91–138 (John Wiley & Sons, Inc, Hoboken, NJ, US, 2002).

31. Gescheider, G. A. in Psychophysics: Method, Theory, and Application 304 (Lawrence Erlbaum Associates Inc, 1985).

32. Fechner, G. T. Elemente der Psychophysik, 1998).

33. Stevens, S. S. & Bartley, S. H. in Handbook of experimental psychology ix, 1436 p. (J. Wiley, New York, 1951).

34. Stevens, S. S. in Psychophysics: Introduction to Its Perceptual, Neutral, and Social Prospects (Wiley, New York, 1975).

35. Short, L. A., Mondloch, C. J., deJong, J. & Chan, H. Evidence for a young adult face bias in accuracy and consensus of age estimates. Br. J. Psychol. 110, 652–669 (2019).

36. Pilz, K. S. & Lou, H. Contextual and own-age effects in age perception. Exp. Brain Res. 240, 2471–2480 (2022).

37. Yoshimura, N., Yonemitsu, F., Sasaki, K. & Yamada, Y. Robustness of the aging effect of smiling against vertical facial orientation. F1000Res 11, 404 (2022).

38. Ganel, T., Chajut, E. & Algom, D. Visual coding for action violates fundamental psychophysical principles. Current Biology 18, R599–R601 (2008).

39. Heath, M., Manzone, J., Khan, M. & Davarpanah Jazi, S. Vision for action and perception elicit dissociable adherence to Weber’s law across a range of ‘graspable’ target objects. Exp. Brain Res. 235, 3003–3012 (2017).

40. Ganel, T., Freud, E. & Meiran, N. Action is immune to the effects of Weber’s law throughout the entire grasping trajectory. Journal of vision 14 (2014).

41. Yarkoni, T. & Westfall, J. Choosing Prediction Over Explanation in Psychology: Lessons From Machine Learning. Perspect. Psychol. Sci. 12, 1100–1122 (2017).

42. Ganel, T. & Goodale, M. A. The effects of smiling on perceived age defy belief. Psychon. Bull. Rev. 25, 612–616 (2018).

43. Ganel, T. Smiling makes you look older. Psychon. Bull. Rev. 22, 1671–1677 (2015).

44. Thorley, C. How old was he? Disguises, age, and race impact upon age estimation accuracy. Appl Cognit Psychol 35, 460–472 (2021).

45. Clifford, C. W. G., Watson, T. L. & White, D. Two sources of bias explain errors in facial age estimation. R. Soc. Open Sci. 5, 180841 (2018).

46. Minear, M. & Park, D. C. A lifespan database of adult facial stimuli. Behav. Res. Methods Instrum. Comput. 36, 630–633 (2004).

47. Zhang, Z., Song, Y. & Qi, H. Age Progression/Regression by Conditional Adversarial Autoencoder, 2017).

48. Moyse, E. & Brédart, S. An own-age bias in age estimation of faces. European Review of Applied Psychology 62, 3–7 (2012).

49. Anastasi, J. S. & Rhodes, M. G. An own-age bias in face recognition for children and older adults. Psychon. Bull. Rev. 12, 1043–1047 (2005).

50. Dibeklioğlu, H., Alnajar, F., Salah, A. A. & Gevers, T. Combining facial dynamics with appearance for age estimation. IEEE Trans. Image Process. 24, 1928–1943 (2015).

51. Moyse, E., Beaufort, A. & Brédart, S. Evidence for an own-age bias in age estimation from voices in older persons. European Journal of Ageing 11, 241–247 (2014).

